# First detection of Zika virus in neotropical primates in Brazil: a possible new reservoir

**DOI:** 10.1101/049395

**Authors:** Silvana Favoretto, Danielle Araújo, Danielle Oliveira, Nayle Duarte, Flavio Mesquita, Paolo Zanotto, Edison Durigon

## Abstract

Samples from sera and oral swabs from fifteen marmosets (Callithrix jacchus) and nine capuchin-monkeys (Sapajus libidinosus) captured in Ceara State in Brazil were tested for Zika virus.  Samples were positive by Real time PCR and sequencing of the amplified product from a capuchin monkey showed 100% similarity to other ZIKV from South America. This is the first report on ZIKV detection among Neotropical primates.

---

In Brazil, the first diagnosed case of Zika virus (ZIKV) in humans was reported in 2015 in Rio Grande do Norte State in the Northeast region, with subsequent reports in 22 States of the Country^1^. We tested samples from sera and oral swabs from fifteen marmosets (*Callithrix jacchus*) and nine capuchin-monkeys (*Sapajus libidinosus*) captured (SISbio license 45196-3) from July to November of 2015 in Ceará State, an epidemic area for ZIKV. Preliminary detection of these samples indicated 29% of positivity (7/24) by Real time PCR^2^ (Cycle Threshold *Ct* average 31,64 to 37,78). We found four positive samples from marmosets and three positive samples from capuchin monkeys. To further validate the Real time PCR we sequenced the amplified product from one capuchin monkey and one marmoset, and found that it had ZIKV with 100% similarity to other ZIKV from South America (Table 1). Primates were captured at several, far apart regions of the State, including coastal, savannah (caatinga biome) and a mountain range with rainforest remnants. It is noteworthy that marmosets were free ranging, co-inhabiting with humans; the capuchins were pets except for one that was kept in a screening center for wild animals in Fortaleza (capital of the State and coastal region). After sample collection, they were tagged with microchips and released back to their original site. Currently, Brazil is facing a massive ZIKV epidemic, associated with severe neuropathy. Three hundred and sixty three cases of microcephaly were notified in Ceará alone in the period from October 2015 to March 2016. Out of these, seventy (19·3%) had laboratory confirmation to related to ZIKV infections^3^. Crucially, cases of microcephaly occurred in municipalities from where viremic monkeys were sampled (Figure 1). This is the first report on ZIKV detection among Neotropical primates, which stands as a caveat for the possibility that they could act as reservoirs, similar to the sylvatic cycle of yellow fever in Brazil^4^.

**Table 1:** Register number, animal, species, Cycle Threshold (*Ct*) and e-value from GenBank of samples from neotropical primates positive for Zika virus in Brazil.

| Number | Animal | Species | *Ct* | E-value (BlastN search) |
| --- | --- | --- | --- | --- |
| bruspCE05_15 | Capuchin monkey | *Sapajus libidinosus* | 37.20 | not sequenced |
| bruspCE08_15 | Capuchin monkey | *Sapajus libidinosus* | 31.34 | 3e -148 |
| bruspCE46_15 | Marmoset | *Callithrix jacchus* | 37.32 | not sequenced |
| bruspCE47_15 | Marmoset | *Callithrix jacchus* | 37.43 | not sequenced |
| bruspCE59_15 | Marmoset | *Callithrix jacchus* | 35.14 | not sequenced |
| bruspCE63_15 | Marmoset | *Callithrix jacchus* | 37.02 | 3e -24 |
| bruspCE68_15 | Capuchin monkey | *Sapajus libidinosus* | 37.73 | not sequenced |

**Figure 1.**
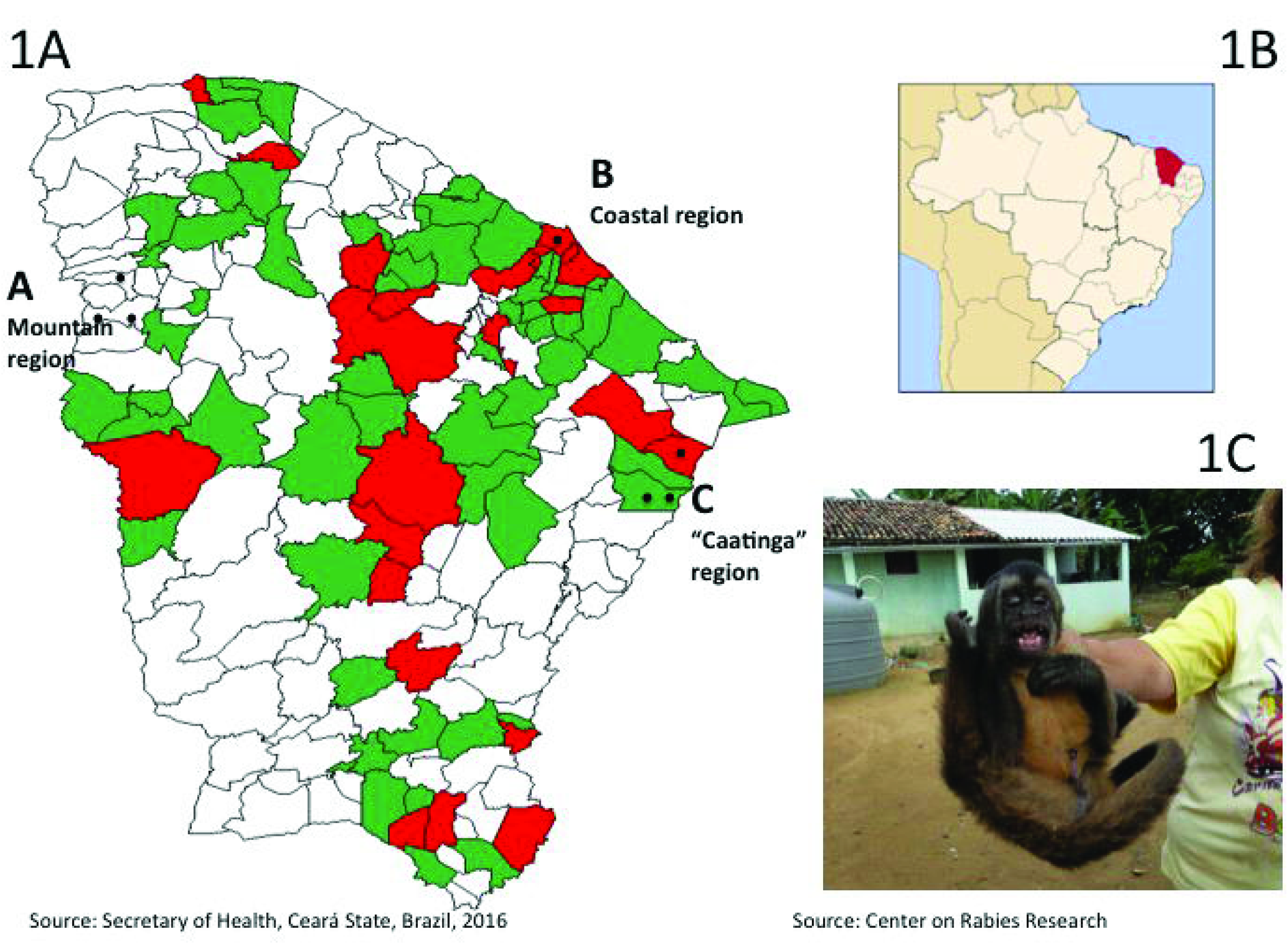
Widespread distribution of ZIKV infected primates in distinct biomes in the Northeast of Brazil. **1A)** Map of Ceará State, Brazil showing cities with notified (green) and confirmed (red) cases of microcephaly and congenital malformations, associated with ZIKV infections. Black dots indicate neotropical primates found to be positive for ZIKV in real time PCR. Two positive marmosets and one positive capuchin monkey was found in the mountain range. One positive capuchin monkey was found in the Coastal region. Two marmosets and one capuchin monkey were found in the savannah (caatinga) region. **1B)** Position of the Ceará State in the Northeast of Brazil. **1C)** A positive pet capuchin monkey with its owner.

This work was funded by FAPESP project Nº 2014/16333-1

